# Decoding in the fourth dimension: Classification of temporal patterns and their generalization across locations

**DOI:** 10.1101/2024.06.19.599699

**Authors:** Alejandro Santos-Mayo, Faith Gilbert, Laura Ahumada, Caitlin Traiser, Hannah Engle, Christian Panitz, Mingzhou Ding, Andreas Keil

## Abstract

Neuroscience research has increasingly used decoding techniques, in which multivariate statistical methods identify patterns in neural data that allow the classification of experimental conditions or participant groups. Typically, the features used for decoding are spatial in nature, including voxel patterns and electrode locations. However, the strength of many neurophysiological recording techniques such as electroencephalography or magnetoencephalography is in their rich temporal, rather than spatial, content. The present report proposes a new decoding method that relies on the time information contained in neural time series. This information is then used in a subsequent step, generalization across location (GAL), which characterizes the relationship between sensor locations based on their ability to cross-decode. Two datasets are used to demonstrate usage of this method, referred to as time-GAL, involving (1) event-related potentials in response to affective pictures and (2) steady-state visual evoked potentials in response to aversively conditioned grating stimuli. In both cases, experimental conditions were successfully decoded based on the temporal features contained in the neural time series. Cross-decoding occurred in regions known to be involved in visual and affective processing. We conclude that the time-GAL approach holds promise for analyzing neural time series from a wide range of paradigms and measurement domains providing an assumption-free method to quantifying differences in temporal patterns of neural information processing and whether these patterns are shared across sensor locations.

**Author summary:** Decoding and classification approaches are widely used in computational biology. In the field of neuroscience, pattern classification approaches typically use spatial information. In many instances, neural time series however are best defined by the their temporal, rather than spatial, features. Here, we propose a novel decoding approach taking advantage of the temporal information. Specifically, we utilize the waveform of neural time series as features for decoding experimental conditions and quantify each decoder’s generalization across locations (GAL) in multi-channel recordings. We illustrate the usage of our open source toolbox using two datasets, showing the sensitivity of the method to systematic condition differences in temporal dynamics along with its ability to capture and quantify spatial dependencies between recording locations.

## Introduction

The development of human neuroimaging tools such as functional magnetic resonance imaging (fMRI) and magneto- and electroencephalography (M/EEG) has transformed the study of human brain function. Traditionally, these measurements have been analyzed using univariate statistics [1], in which a linear statistical model is applied to a given time point, voxel, or electrode. Because of the univariate nature of this approach, it disregards information regarding spatial or temporal patterns [2]. Thus, multivariate pattern analysis (MVPA) has emerged as an alternative tool for discriminating the neural activity patterns related to a given stimulus, experimental condition, or participant [3,4].

Multivariate decoding techniques capitalize on the information found in neural patterns to characterize the multifaceted properties that define a given brain process [5]. Also known as decoding methods, these techniques were originally introduced for fMRI data [4,6], and their use has steadily increased in recent years, becoming a standard procedure in this field [5,7]. Meanwhile, in the field of M/EEG research, decoding approaches are also increasingly used [8] with several standard procedures emerging [9–11]. Generally, these standard procedures use spatial information to decode the underlying neural activity patterns, paralleling approaches used with fMRI data (see Figure 1 left). However, the resolution of spatial information is typically reduced for electrophysiological tools like EEG in comparison with fMRI. It is widely accepted that the primary strength of EEG and MEG recordings lies in their rich temporal information, rather than in their spatial distribution. Specifically, human electrophysiology reflects the time series of neural population activity arising from post-synaptic events, subject to volume conduction and other distortions when measured from the scalp. The resulting waveforms contain valuable information about the neural response and/or oscillatory rhythms related to a specific experimental condition or stimulus [12].

**Figure 1:**
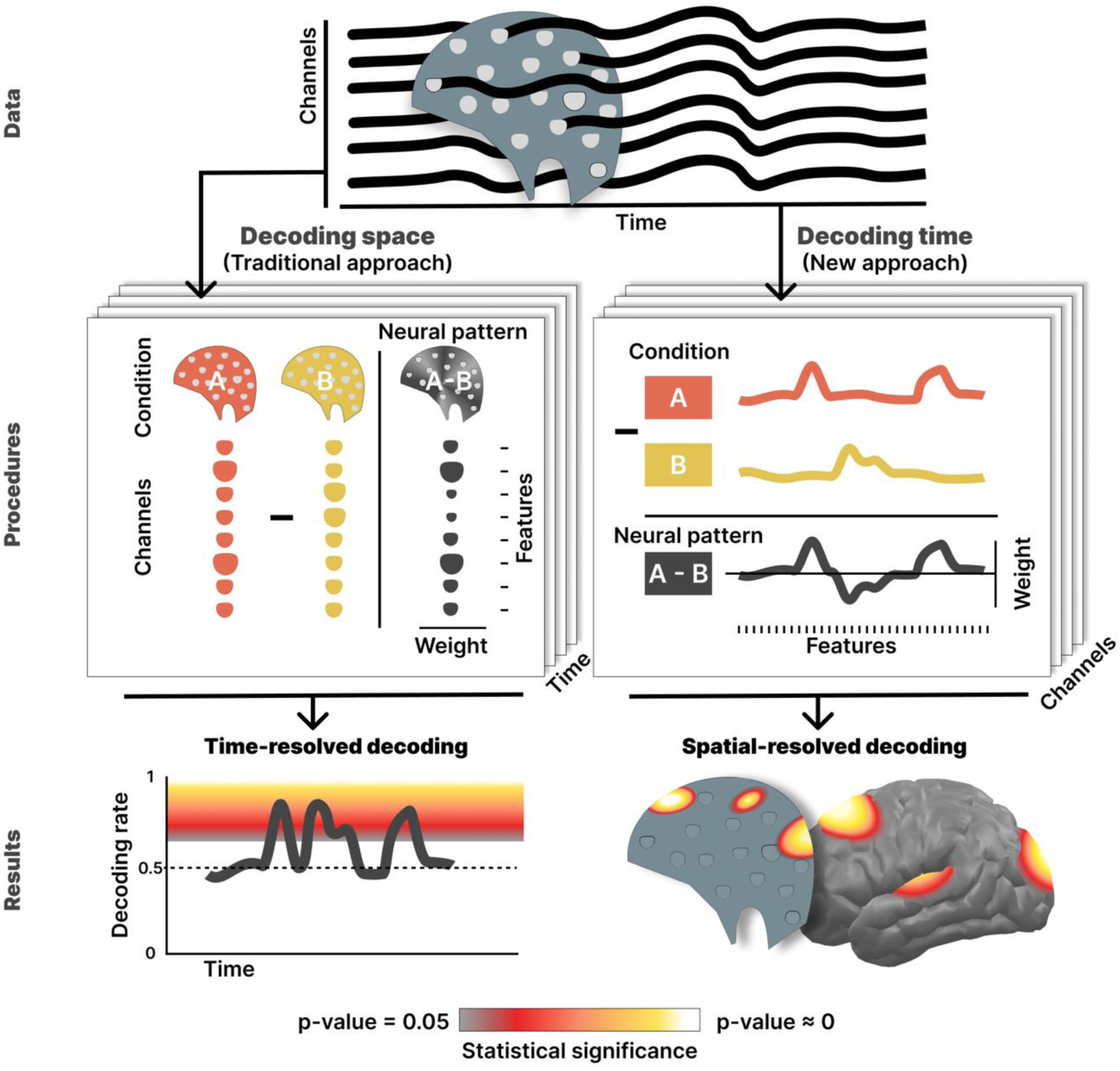
Flowchart describing how MVPA-based decoding techniques may be carried out following the traditional approach (**left**) or the time-GAL approach proposed here (**right**). Data from electrophysiological recordings (**top**) offer both spatial and temporal information in the form of channels and time series, respectively, which can be used for decoding. Thus, each procedure (**middle panles**) leverages one of the dimensions using its data to fit a classifier, repeating this operation over the other dimension to examine generalization. Specifically, the traditional approach employs channels or sensors as features; the time-GAL approach uses temporal features. Finally, the results (**bottom**) for both the traditional (space-based) and the time-GAL method show the decoding performance across time or locations, respectively.

In the present study, we apply MVPA algorithms to the temporal dimension of EEG signals, taking advantage of the high resolution of neural time series data in this dimension. In this new method, we use all time points of each trial time series as features that enter a multivariate classifier model (see Figure 1, right), in contrast to spatial information (recording sites) which has been used in the literature previously. Then, we repeat this procedure for each recording location (e.g. electrodes in the case of EEG) to obtain a spatially resolved depiction of scalp regions in which the temporal dynamics related to a condition or stimulus contain overlapping decodable information based on temporal features. In a second step, we adapt a method proposed by King and Dehane [13] called generalization across time (GAT), in which a classifier algorithm is trained using spatial information for one (training) time point and then tested across a range of different time points. This procedure allows researchers to characterize the temporal stability of neural patterns related to specific conditions or stimuli. The present approach, using the temporal waveform as a feature vector for classification allows us to quantify the extent to which different recording sites share decodable information. This is implemented as a generalization across location (GAL), in which a temporal decoder trained at one location is applied to other locations. By mapping the cross-decoding accuracy across locations, the time-GAL method offers a multivariate approach to defining the unique topographic patterns in which temporal dynamics are related to a specific stimulus or experimental condition. Finally, another important contribution of the current work consists of the combination of not only spatial, but also temporal information to the decoding output. Classifier models are considered backward models as they extract the information about experimental manipulations from the brain’s responses to these manipulations. On the other hand, forward models describe the process in which experimental manipulations and brain biophysics lead to the recorded neurophysiological data. The exact weights of a classifier-based backward model can only be extracted using a corresponding linear forward model [14]. Therefore, we implemented a simple linear forward model based on correlations of the data with the labels to visualize the dynamics of time-GAL decoding and cross-decoding patterns.

In the present study, we demonstrate the usage of the time-GAL approach for analyzing neural time series data, using two EEG experiments. The experiments involved (1) affective picture viewing and (2) aversive generalization conditioning.

## Methods

### Datasets

To provide a demonstration of the time-GAL procedure, two datasets were used, involving two types of electrophysiological response: event related potentials (ERP) and steady-state visual evoked potentials (ssVEP). ERPs are typically generated by the repeated presentation of transient stimuli while the EEG is recorded, often at temporal intervals of several seconds. Stimulus-locked EEG segments obtained for each presentation are then averaged, resulting in a prototypical voltage time course [12,15]. By contrast, ssVEPs result from rhythmic presentation of visual stimuli (e.g. regularly flickering light), driving an oscillatory neural response at the same temporal frequency as the driving stimulus [16,17].

Data set 1 comes from a picture viewing task in which ERPs were measured from 39 participants (27 women; average age 19.6 years) while passively watching pleasant, neutral, and unpleasant affective pictures from the international affective picture system (IAPS; [18]. The present demonstration focuses on two conditions, pleasant and unpleasant picture viewing. These conditions were selected because they each prompt robust ERP signals but tend to show only small condition differences when compared using univariate methods [19]. 20 pleasant and 20 unpleasant pictures were shown twice each, for a duration of 1 second at the central portion of the screen, resulting in a total of 80 trials per participant, with 40 trials in the pleasant and 40 in the unpleasant condition.

Data set 2 comes from a fear conditioning paradigm in which neutral cues were flickered at 15 Hz, prompting an oscillatory SSVEP response, recorded in 31 participants (22 women; average age 19.23 years). Visual cues consisted of 4 high-contrast circular sinusoidal gratings (orientations: 15°, 35°, 55° and 75°) filtered with a Gaussian envelope (Gabor patches). Each stimulus was presented against a dark gray background in a flickering (15 Hz) mode for 3 seconds. The fear generalization protocol [20] involved two phases: First, in the habituation phase, each of the 4 visual cues was presented 30 times in random order. Later, during the fear acquisition phase, the 4 visual cues were again shown, but one of them (CS+, 15° or 75° randomly assigned per subject) was followed by an aversive loud (88 or 91 SPL dB) white noise (the unconditioned stimulus, US) that started 2 seconds after the onset of the CS for a duration of 1 second, thus co-terminating with the CS. After acquisition, the CS+ orientation typically acquires aversive properties, while the remaining 3 orientations served as generalization stimuli (GS) with increasing dissimilarity relative to the CS+ based on their orientation. Thus, GS1 was the closest stimulus to CS+ in orientation and GS3 was the most dissimilar one.

Procedures of both experiments were approved by the University of Florida institutional review board and informed consent was signed by all participants in accordance with the Declaration of Helsinki.

### EEG data acquisition and preprocessing

EEG signals from both datasets were collected using an Electrical Geodesics (EGI, Eugene, OR) system with a 128-channel HydroCel electrode net, and continuously digitized at 500 Hz, referenced to the vertex sensor (Cz). Electrode impedances were kept below 50 kΩ. Online (low-pass filter, 3 dB point at 170 Hz; high-pass filter, 3 dB point at 0.05 Hz) and offline (low-pass filter, 40 Hz, 18th order, 3 dB point at 0.05 Hz) Butterworth filters were applied to the data. Then, the signal was segmented in epochs including 0.2 s pre-stimulus and 1 s post-stimulus for the ERP and 1 s pre-stimulus, and 2 s post-stimulus for the ssVEP datasets. Statistical Correction of Artifacts in Dense Array Studies (SCADS) procedure [21] was employed for artifact rejection of bad channels and epochs. Finally, all trials were individually baseline corrected by subtracting the voltage activity average in the pre-stimulus segment of the same trial and ordered in categories pleasant (1037 trials across all participants) and unpleasant (1133 trials) for the ERP dataset and CS+ (548 trials), GS1 (570 trials), GS2 (571 trials), GS3 (578 trials) for habituation and acquisition phases of the ssVEP dataset.

### Current source density (CSD)

EEG recordings are characterized by pronounced spatial smoothing and distortion as signals travel from the brain to the extracranial sensor locations, caused by a number of biophysical processes including volume conduction and spatial lowpass filtering. The reduction of shared information is crucial in terms of assessing the spatial generalization of time-based decoders [22]. Here we applied the current source density (CSD) transformation, an established algorithm for diminishing spatial smearing in the topographical distribution of EEG signals [23]. Specifically, we used the implementation by Junghöfer et al [21] setting the smoothing (regularization) parameter λ to a value of 2.

### Decoding analysis for brain response generalization across spatial and time dimensions

#### Electrophysiological waveform-based classification

In the current report, we first apply (train) a linear discriminant decoder to discriminate two conditions based on temporal features of the electrophysiological signal. Then, in a second step, we quantify the generalization of the temporal decoders across channels. This approach is similar to the generalization across time (GAT) method [13], where a spatial decoder is implemented (trained) to discriminate conditions within one or more time points, followed by its application (testing) across a range of different time points. Although GAT sheds light on the decoding time course, the topographic information and the relation between different sensor locations cannot be seen. This report introduces a new method, the generalization across locations (GAL) approach, to address this gap. In contrast to the GAT method, the time-GAL algorithm takes advantage of the temporal information, i.e. the signal waveform, to train a decoder for each channel. Each channel’s decoder is then tested at all other channel locations, a process known as cross-decoding. Because this decoding approach aims to infer neural processes from data, it can be considered a backward model [14]. In contrast, a forward model is a model that describes how brain activity results in neural data as collected for example by EEG. Because there is a corresponding forward model to each backward model (Haufe et al. [14]), we can employ a forward modeling approach to obtain the information regarding when each topographic channel reflects condition differences.

#### Backward model of the generalization across location

For the GAL analysis, we use data from two conditions, referred to as A and B. In our implementation, each data set consisted of three-dimensional arrays (channels by time points by trials) for each condition and each subject (see Figure 2 top left). For one of *k* channels, we fed the classifier model all time points as features and all single trials from all *n* participants except from one participant (leave-one-out training/testing), as observations. Because the classifier operates at the single trial level, many observations were considered for the training phase of the classifier model, creating pressure on computing resources. Therefore, we compared the performance and efficiency of a linear discriminant analysis (LDA) with a support vector machine (SVM). LDA is proven to be suitable for neural decoding [25], although SVM has been more widely used in EEG data [10,26] for averaged ERPs. Similar to Grootswagers et al. [8], we found comparable decoding rates from both types of models but a significant reduction of time consumption from the LDA algorithm. Because we perform a substantial amount of classifier training and we seek a computationally efficient methodology, we selected LDA as our classifier model to discriminate training trials as coming from conditions A or B. Models were trained using the LDA function built in MATLAB (2023b). Even when the number of trials were unbalanced, all single trials from conditions A and B were employed to avoid randomness in any aleatory selection of trials. Priors were set accordingly. Information contained at single-trials is noisier than evoked information from trial-averages (e.g., ERPs), but single-trial time courses include the non-phase-locked temporal features of the signal, often referred to as “induced”, instead of “evoked” responses [12]. More information about the difference between applying the GAL methodology to single-trial or averaged is given in S1 Appendix.

In the second step of the time-GAL method, we applied the classifier model weights trained at a given channel to quantify its decoding performance with data from all *k* channels using the single trials of the remaining subjects as the test data set. Thus, we required the decoder to use feature weights that were applicable across participants. The GAL test phase provides a classification of trials into A and B that quantifies the accuracy of the model using the decoding accuracy (correct A and B classification) as an index. This procedure was repeated *k* times, training the classifier model with all the remaining channels and testing their weights with all the *k* channels to fill out the decoding accuracy in a generalization matrix of training by testing axes with dimensions *k* by *k* channels (see Figure 2 middle right). Finally, this analysis was computed again for each subject, using the rest of the *n* - 1 participants as training dataset in a leave-one-out approach. Thus, this approach to cross-validation avoids arbitrary decisions (e.g. regarding the number of k-folds) in the training/test data set selection. Finally, this methodology provides us with *n* generalization matrices of channel-by-channel dimension, where *n* corresponds to the number of subjects. This distribution is employed later in the following statistical analyses. Conceptually, if a time-based decoder trained at location *k*_i_ also classifies the data at location *k*_j_, then we interpret this as evidence that the temporal dynamics at *k*_i_ and *k*_j_ contain share decodable information about the conditions—a non-parametric way of describing spatial dependency or connectivity.

**Figure 2:**
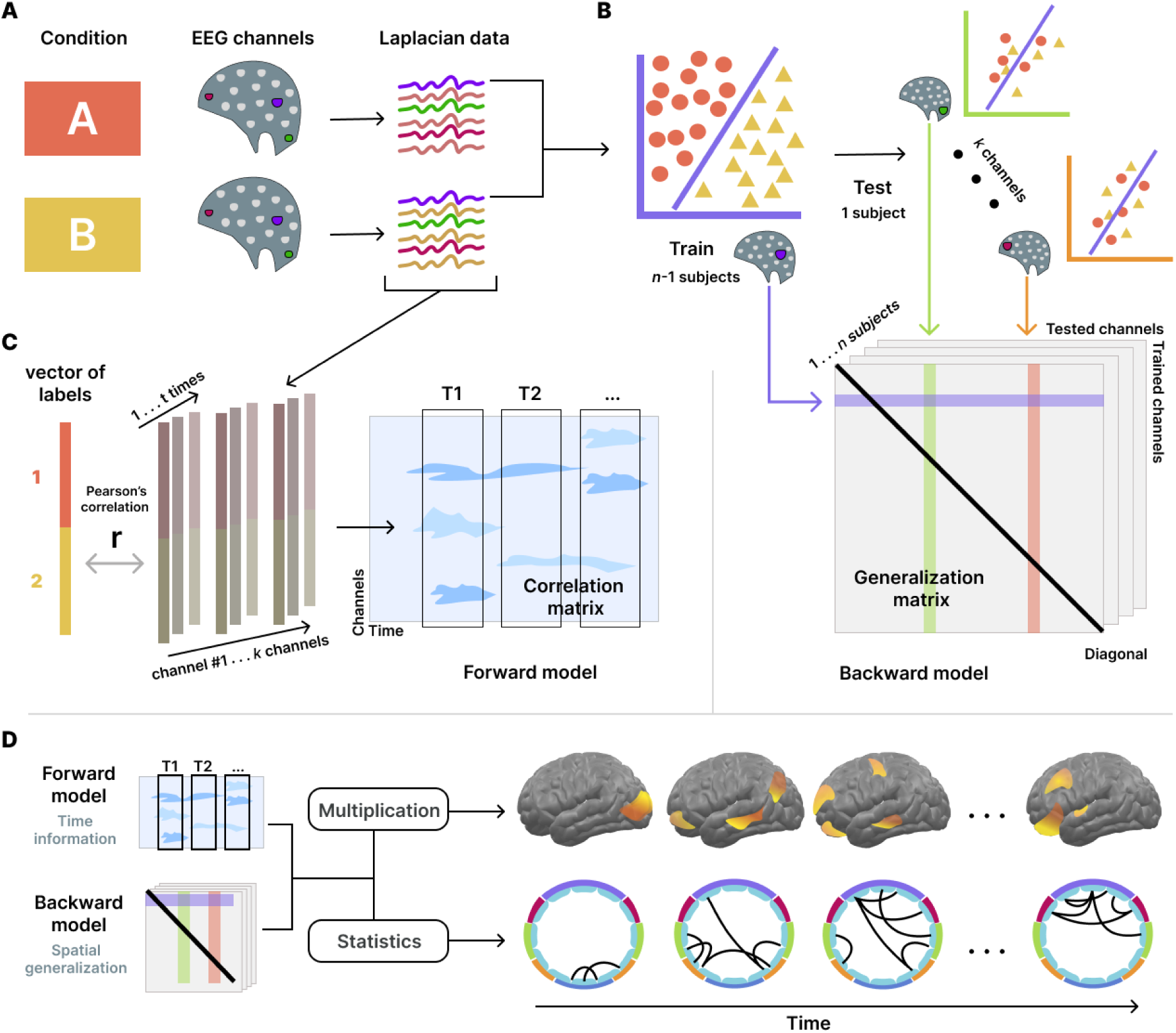
Flowchart describing the decoding methodology based on the temporal and spatial information. **A**: Laplacian EEG data from conditions A and B are used to train one classifier per channel using time points as features. **B**: Next, the generalization matrix (backward model) is calculated by testing every classifier model with data from each of the remaining channels (cross-decoding). **C**: Then, the correlation matrix (forward model) is calculated using Pearson’s correlations between the A/B labels and EEG data for each channel and time. **D**: Finally, the combination of both backward and forward models containing the spatial and temporal information, respectively, allows us to visualize the GAL cross-decoding patterns in the brain. We interpret these as similarity or connectivity matrices, akin to correlation-based connectivity matrices used in EEG and fMRI research.

#### Forward model of the temporal contribution

The GAL method allows us to inspect the spatial distribution of the decoding performance between A and B conditions, based on the electrophysiological waveform, or time information. However, this approach offers only a static picture of the connectivity matrix. In the GAL framework, a “connection” indicates that two channels share decodable information. Therefore, it is helpful to visualize what time points in the waveform are most strongly related to the discrimination of conditions. As stated above, it is possible to estimate a forward model to obtain the weights of the LDA features [14]. We can address this estimation by measuring the relations between the condition labels and the Laplacian EEG data. To this end, the Pearson’s correlation coefficient between the true vector of A and B labels and all subjects’ trials of electrophysiological activity was calculated for each time point and channel resulting in a channel by time matrix of correlations (see Figure 2 middle left).

#### Combining decoding topography with the time domain

While the backward model provides spatial information, we use the forward model to visualize the temporal progression of condition differences at the single trial level. Specifically, we combine the information from both models to obtain a temporally weighted representation of the decoding patterns. First, we were interested in a brain topographic visualization of the main contributor areas to the decoding analysis. To obtain this visualization, we used the diagonal vector from the average of the *n* generalization matrices, i.e., the decoding accuracy of each channel’s LDA, across participants. This information served to project the decoding rates of the *k* channels into the source space. Then, the decoding-weighted forward model is obtained by timepoint-wise multiplication of the overall decoding vector of *k* spatial values with the absolute values of the *k* by *time* correlation matrix. The absolute value of the correlation matrix is used because it indicates the contribution of a given channel and time point to the discrimination between conditions A and B, regardless of the sign. This combination of the decoding vector (spatial information) and correlation matrix (temporal information) results in another *k* by *time* matrix, which reflects the time-weighted spatial contribution to the decoding. Thus, we obtain one topography that represents the spread of decodable information across the brain surface for every *time* step. EMEGS 2.8 [27] was used to project this information onto a standard cortical surface, for visualization.

#### Statistical extraction of GAL connectivity patterns over time

For the statistical analysis of GAL data, a mass univariate approach was chosen. Once the *n* generalization matrices were computed, Student’s T test against the decoding chance rate were computed for each element in both directions, i.e., above and under the chance level. Since we only have conditions A and B, the decoding chance rate of the classifier is 0.5. The diagonal of the GAL matrix shows the decoding accuracy for the test dataset of the same channel used for classifier training, and thus its values are not expected to be smaller than chance, i.e. 0.5. However, the values for the rest of the generalization matrix, in which test and training datasets came from different channels may well show values below chance level. This would indicate that the data from one location are systematically located on the other side of the hyperplane spanned by the LDA trained on another location. This may be interpreted as those locations showing orthogonal neural dynamics, more like the other label, resulting in significantly below-chance decoding. Therefore, we used the contrast T test in both directions showing not only those positive connections between regions, but also negative connections (below chance cross-decoding). Each of these analyses resulted in one matrix of channel-by-channel dimensions showing a large number of comparisons. Since connections between different channel locations cannot be considered spatially near, i.e. neighbors, their independence can be assumed, and Bonferroni correction was applied to address the multiple-comparison problem. Also, to obtain only the most meaningful generalization connections, a corrected alpha (α) was utilized (see results). Therefore, only those connections presenting a p-value less than 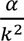 where α represents the desired alpha level, were considered statistically significant and used for the description of the connectivity (GAL) pattern.

For visualization, in a final step, we combined the statistically significant GAL results with the statistically significant temporal correlations to illustrate the time-GAL dynamics. Specifically, we only considered the Pearson’s correlation significance of those channels that showed statistically significant generalization connections (see above). We used a threshold of α = 0.01 to identify temporal correlations (contributions) of each location that were relevant for visualization. This alpha value was not Bonferroni corrected because it was only used as a threshold for visualization. This step allowed us to visualize the relationship between areas throughout the temporal progression of the neural information processing. Thus, we can inspect how different networks or areas are implicated in different time steps, providing a descriptive video of the cognitive processing instead of a static picture of all the GAL pattern.

The methodology described above is implemented in an open-source MATLAB toolbox. This toolbox can be found in https://github.com/csea-lab/time-GAL and example data may be downloaded at https://osf.io/q56ns. Also, scripts for calculating the forward model through Pearson’s correlation and for combining its temporal information with GAL outputs are included.

## Results

The time-GAL decoding methodology uses a classification approach to quantify decodable temporal information in neural dynamics, including how this information generalizes across locations. In the following, we first illustrate how the time-GAL methodology captures information contained in the ERP response to transient visual stimuli. Later, we examine the application of time-GAL to ssVEP responses.

### Time-GAL patterns on ERP: Affective pictures presentation

The first dataset consists of CSD-transformed EEG recordings obtained during the presentation of pleasant and unpleasant pictures from a standardized picture database, in a within-participants design. Given this paradigm, we used time-GAL to perform pairwise decoding of the two conditions (pleasant vs unpleasant). The GAL approach was computed and a Bonferroni-corrected α of 0.05 was used as the statistical threshold for the generalization connectivity patterns. Results are shown in Figure 3.

**Figure 3.**
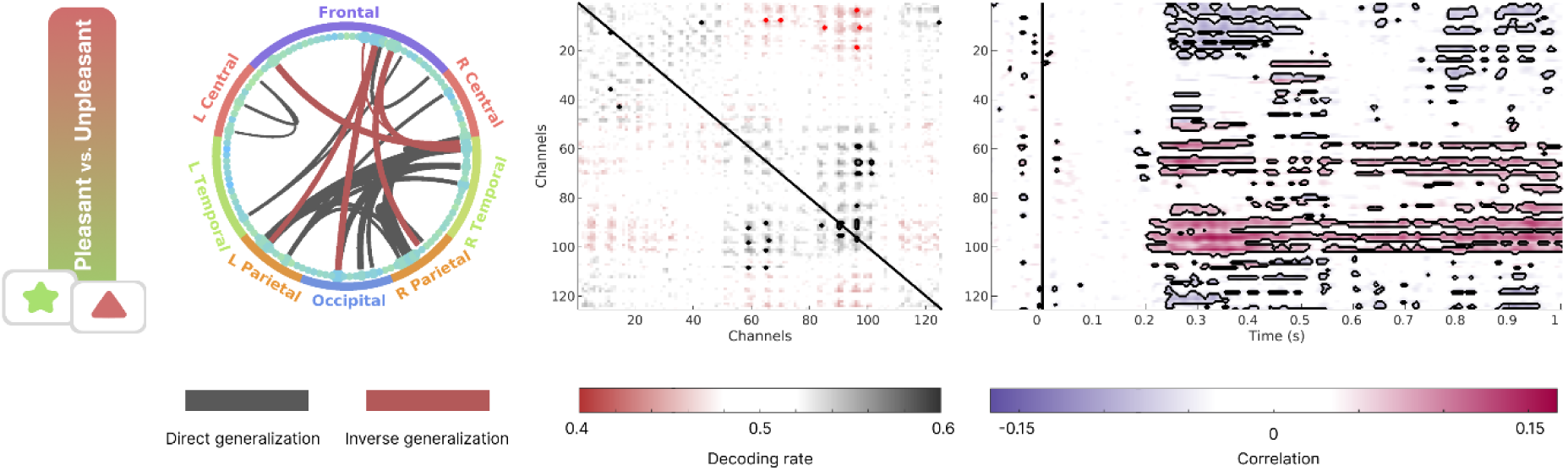
Generalization connectivity patterns and temporal distributions for the affective picture viewing experiment. **Left**: Connectivity graph showing the significant generalization connections between the CSD channels. Black lines refer to connections in which both channels share decodable temporal patterns, while red lines show the opposite pattern (significant below chance decoding). **Middle**: Generalization connectivity matrix showing decoding results (accuracy). The Y axis corresponds to the channel data employed to train the model, and the X axis shows the channel data used for the generalization test. Black and red contours indicate the positive and negative generalization connections, respectively, after a Bonferroni-corrected statistical (α = 0.05) comparison against chance (50%). **Right**: Temporal correlation between labels of condition and data showing the positive or negative weights of the decoding model. Black contours indicate the time points whose p-value is below 0.05 for each channel.

Several posterior channels showed significant decoding based on their time dynamics. Similarly, significant generalization connections were more concentrated among posterior areas. It is worth noting that inverse connections (below chance decoding) were observed between anterior and posterior regions, likely representing opposite voltage of dipolar fields when measured at opposite sides of the head. Latencies of correlation between condition labels and channel and time points indicate the emergence of decodable information around 250 ms after trial onset.

Next, we combined the spatial information of the generalization response across the brain with the temporal weights of the model for each channel. Specifically, we multiplied the diagonal of the generalization connectivity matrix with the Pearson’s correlation matrix of time weights. Additionally, we determined which generalization connections and temporal weights were statistically significant. As shown in Figure 4, both sources of information shed light on cortical surface regions that contain decodable information (brain surface plots, bottom panels) and how they share decodable information (circular graphs).

**Figure 4.**
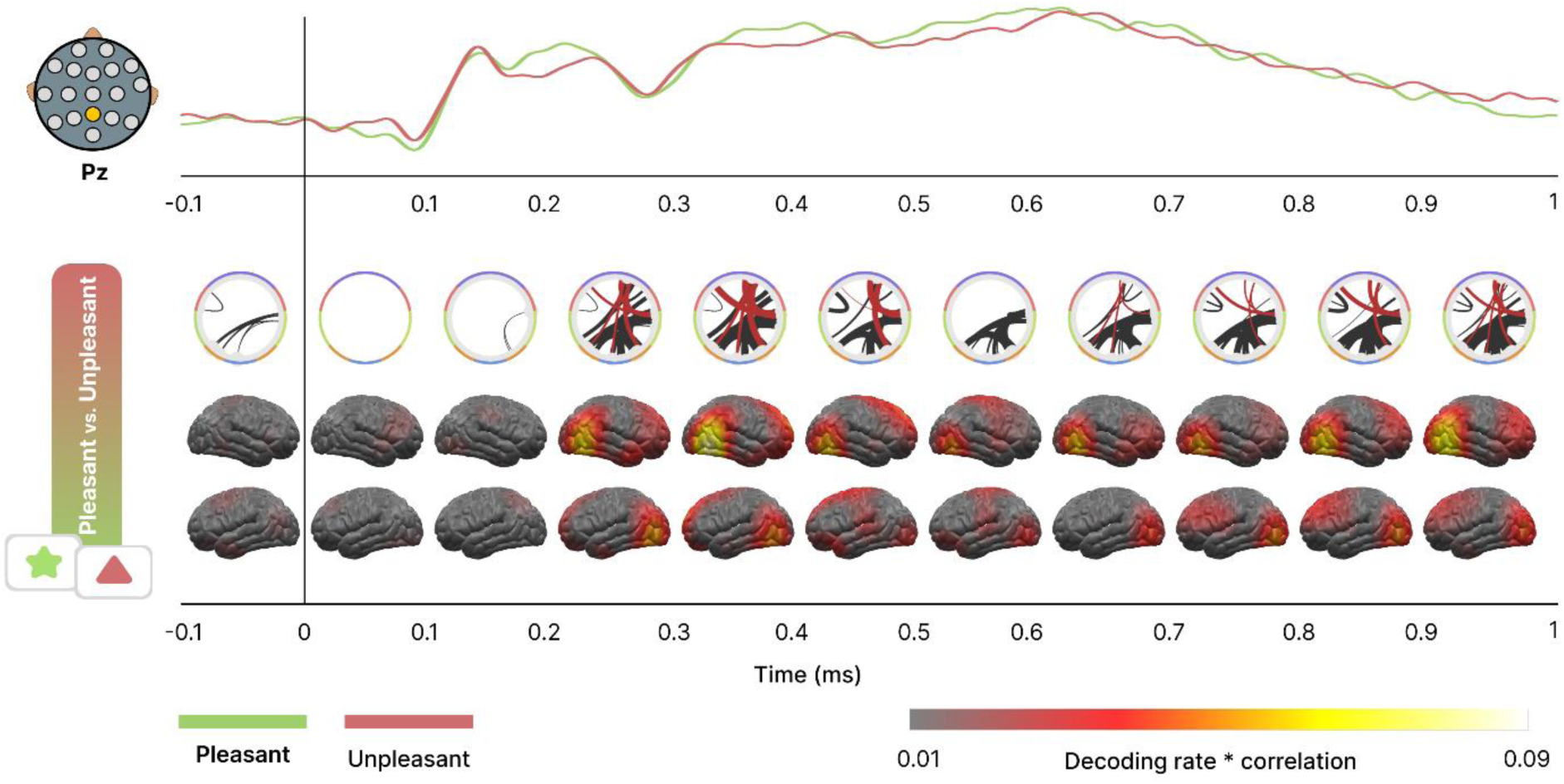
Time course of emotional content decoding and GAL connectivity patterns during affective picture viewing. **Top**: Electrophysiological ERP response to the presentation of different emotional content obtained at the parietal central (Pz) electrode for the three conditions (pleasant, neutral and unpleasant). **Middle**: Circular graphs show the GAL connectivity pattern at each time step for each pairwise condition. The topographies (**Bottom** panel) indicate the multiplication of the decoding rate with the time-varying correlation, showing changes in cross-decoding across the surface of the brain as a function of time.

The time course analysis showed strong inverse (below-chance) decoding between posterior and anterior regions and several direct (positive) generalization connections between right occipital-temporal and parietal zones. The fronto-occipital connectivity pattern was present from 0.2 s to 0.5 s and at the end of the trial from 0.7 s to 1 s, while right occipito-temporal generalization persisted from 0.2 s to the end of the trial. As discussed above, the inverse direction of the GAL results in the antero-posterior connections likely indicates that the voltage patterns found in the occipital and parietal areas are inverted at frontal sensors prompting below chance decoding accuracy. By contrast, the right parietal time-GAL patterns are consistent with studies of the late positive potential as mentioned below in the discussion section.

### Time-GAL patterns in ssVEP signals: Aversive conditioning of oriented Gabor patches

Next, we applied the timed-GAL approach to ssVEP data from an experiment using flickering visual stimuli. Given the high signal to noise ratio of ssVEPs, we employed a more conservative Bonferroni-corrected α of 0.001, to plot only the most salient generalization connections in the GAL analysis. Results presented below in Figure 5 use data from the acquisition phase of the fear conditioning task, meaning that the CS+ stimulus (a grayscale grating defined by its orientation) became a threat cue due to consistent pairing with an aversive white noise (US).

**Figure 5.**
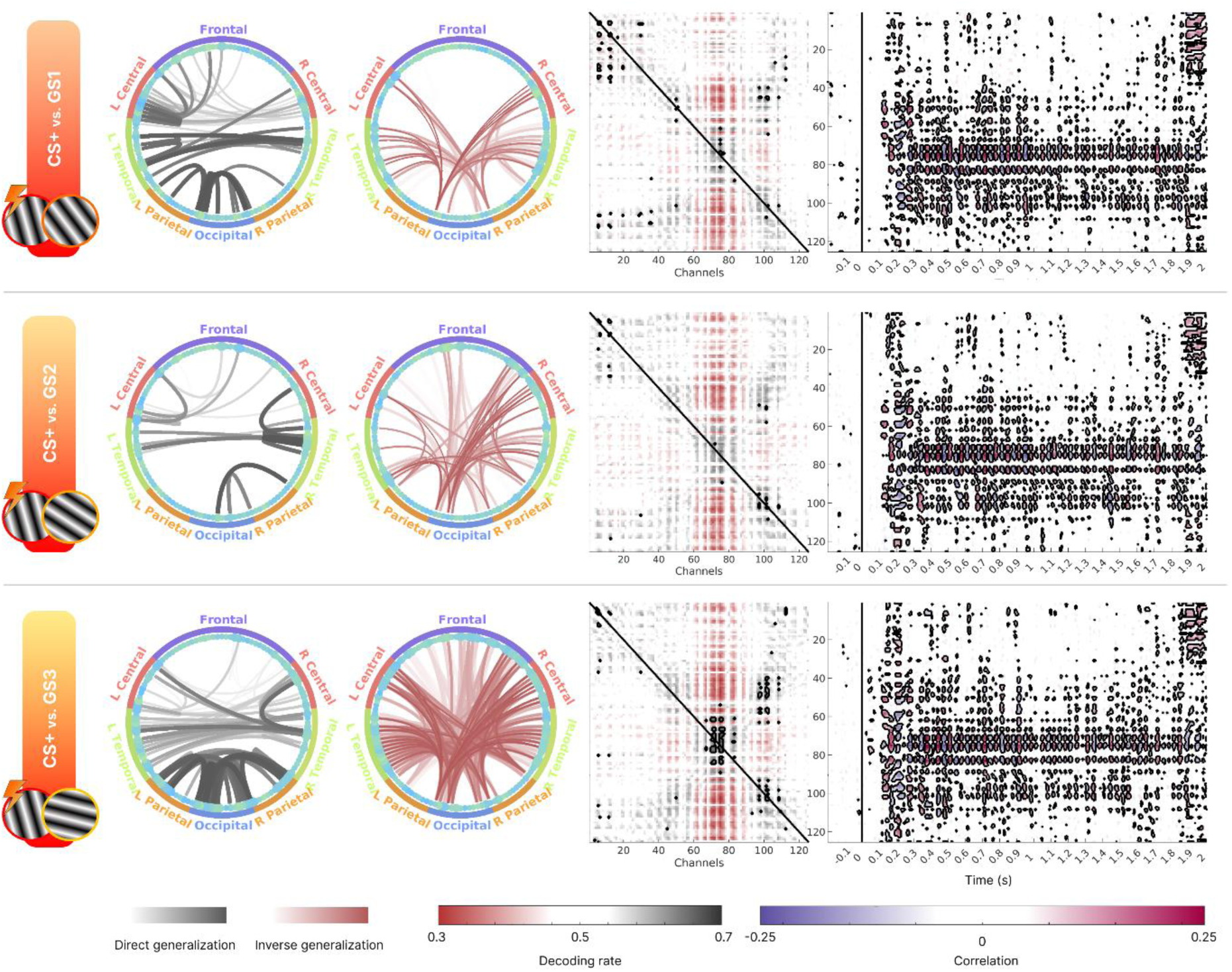
Generalization connectivity patterns during fear conditioning. **Left**: Circular graphs showing the statistically significant generalization connections between 125 CSD channels. Black lines one refers to those connections where both channels show similar decoding temporal patterns, while graph filled with red lines indicate channels showing inversely similar decoding patterns. **Center**: Generalization connectivity matrix showing decoding results (accuracy). Y axis corresponds with the channel data employed to train the model, while X axis refers to the channel data used in the test phase. Black and red contours indicate those direct and inverse generalization connections, respectively, after a Bonferroni-corrected (α = 0.001) statistical comparison with chance level (0.5). **Right**: Temporal correlation between labels of condition and data showing the positive or negative weights of the decoding model. Black contours indicate those time points whose p-value is below 0.01 for each channel.

In terms of direct generalization connections (black lines in Figure 5), the first pairwise comparison, CS+ vs. GS1, shows shared decodable information across frontal and central areas along with the expected occipital areas. Interestingly, direct connections were also found between occipital and frontal areas. The next pairwise comparison, CS+ vs. GS2, showed less frontal generalization and general generalization between temporal and central areas. Finally, the CS+ vs. GS3 comparisons, in which GS and CS+ differed the most, showed strong posterior (parietal and occipital areas) generalization connectivity. Regarding the inverse generalization (red lines in Figure 5), a pronounced generalization was observed, driven by occipital areas, as expected when using ssVEPs. In the first case, the comparison between fear cue and the most similar oriented mainly shows an occipito-temporal connection. However, in the case of the GS2, the inverse generalization from the occipital region reaches all the remaining areas. This spread out of generalization appears more potentiated in the case of GS3, remarkably in the frontal areas. Therefore, the results of the inverse GAL point to an increase and spread out of inverse generalization as a function of dissimilarity between the fear cue and the GS. Thereby, differences between conditions in visual areas would be expressed in the opposite direction along the brain and interestingly at the frontal areas.

Taken together, results indicate distinctive involvement and behavior of anterior and posterior areas directly related with the similarity of the visual object with the threat cue. Finally, compared to the above ERP analysis, the ssVEP shows a fluctuating temporal correlation (Figure 5 right) reflecting the oscillation of the ssVEP between positive and negative voltage. This oscillation in weights for the decoding model was expected since wave-form of the trial, i.e. time dimension, was used as observation for training the model and it corresponds with an ssVEP response. In the case of CS+ vs. GS, all the three pairwise showed a similar temporal weight pattern with a preponderance of visual areas and frontal areas before US onset. We combined the weights with the GAL information as shown in Figure 6.

**Figure 6.**
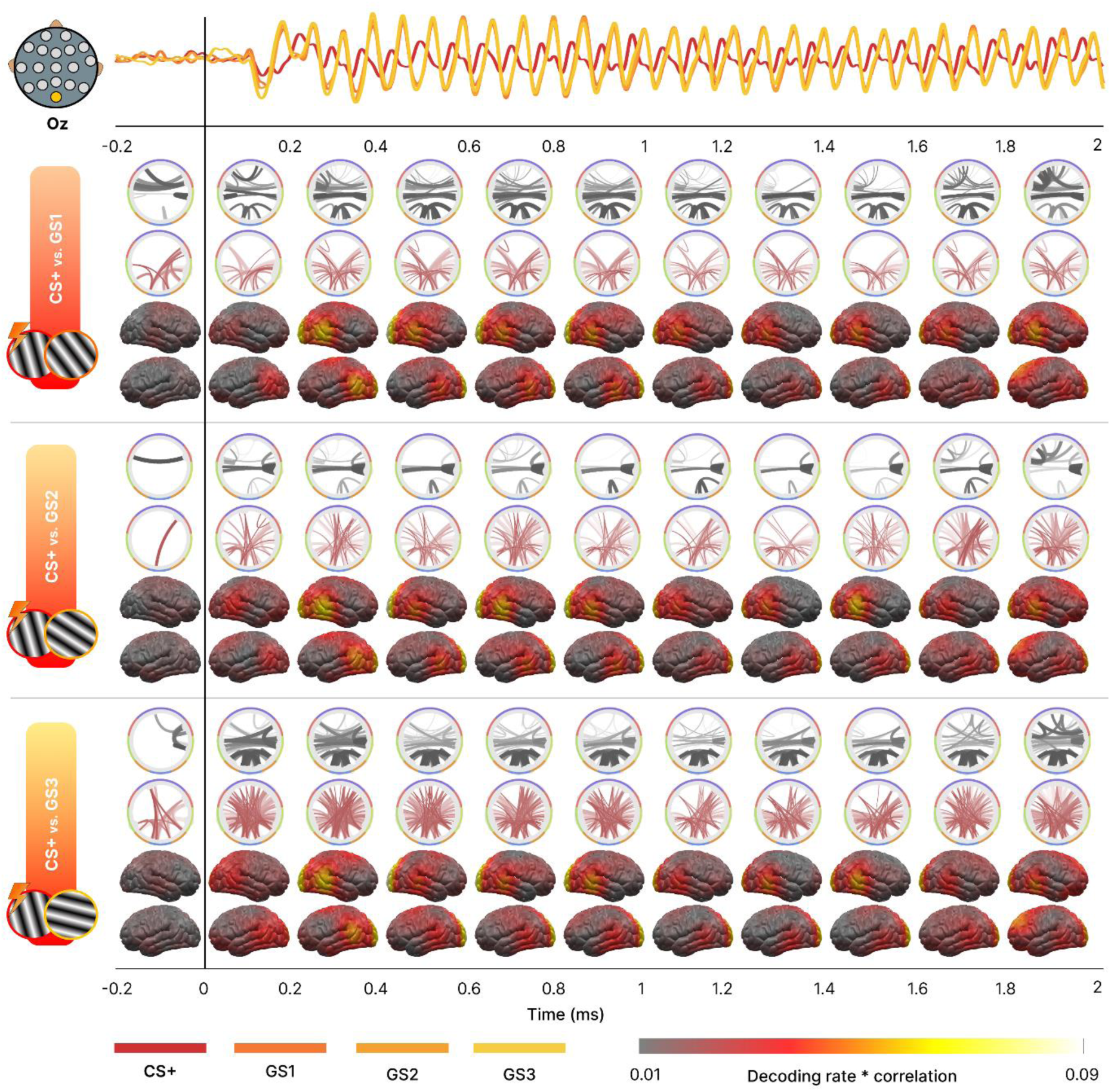
Time course of emotional content decoding and GAL connectivity patterns in fear conditioning. **Top**: Electrophysiological ssVEP response to the presentation of relevant or irrelevant cues obtained at the Occipital central (Oz) electrode for the four conditions (CS+, GS1, GS2, GS3). **Middle**: Circular graphs show the GAL connectivity pattern at each time step for each pairwise condition. For clarity, direct (black lines) and inverse (red lines) generalization across the brain were individually displayed. Brain’s projections indicate the convolution of decoding rate by time correlation describing the fluctuation of the fear discrimination process during the trial over the cortex.

Here, we found that the above-mentioned anterior and posterior modulations take place at the beginning of the trial, after stimulus presentation, towards the end of the trial, and just before onset of the aversive noise (US). Specifically, the projection of decoding accuracy at the source space (see Figure 6) showed a strong involvement of early visual cortex at the beginning of the trial (from 0.2 s). Furthermore, the last 200 milliseconds prior to the onset of the US present a spatial shift in the discrimination between fear relevant or irrelevant cues by recruiting premotor and dorsal regions of the frontal lobe. These occipital and frontal areas implicated in the fear decoding revealed the same latencies and topography for all the three pairwise CS+ vs GS, indicating a similar cognitive processing during the task.

The time-GAL connectivity pattern showed the differences between the three GS responses. Higher direct generalization (black lines in Figure 6) is present in frontal areas of the CS+ vs. GS1 pairwise compared to the inverse generalization (red lines), while the CS+ vs. GS3 pairwise showed the contrary pattern with less direct generalization and a massive frontal involvement in inverse generalization. This frontal modulation was already established, but this final time-GAL step showed that it is most pronounced at the beginning and end of the trial. On the other hand, higher generalization in both direct and inverse direction of the visual and posterior tiers was found for the CS+ vs. GS3 compared with the CS+ vs. GS1 pairwise, indicating the contribution of this regions to discriminate better between different orientations. This posterior differences in generalization persisted throughout the entire trial. Taken together, results of the generalization connectivity matrices showed that the decoding itself (i.e. diagonal of the matrix) did not differ between the three comparisons made here, but the generalization across locations did.

As a control, we repeated the same analysis in the habituation phase of the fear conditioning paradigm. This phase is characterized by the absence of the aversive white noise, i.e. the CS+ orientation is not paired with the US. Therefore, no threat is associated with any of the oriented Gabor patches.

Interestingly, decoding-based GAL analysis conducted on the habituation phase did not reveal any statistically significant decoding (diagonal) or connection (triangles) from the entire generalization connectivity matrix (p > Bonferroni-corrected 0.01). This lack of discriminability between oriented patches highlights the specificity of the effects observed during acquisition.

An alternative way of quantifying and illustrating time-GAL information is shown in Figure 7. Time-GAL information is readily reduced by organizing the density of connections between regions as presented in the middle column of Figure 7. The resulting matrices show the total amount of connections, both direct and inverse, between two areas. The right column of Figure 7 shows differences involving higher connectivity (red squares) in channels of central areas when comparing CS+ vs GS1, compared to other comparisons, while CS+ vs. GS3 showed more connections (blue squares) in temporal and posterior areas.

**Figure 7.**
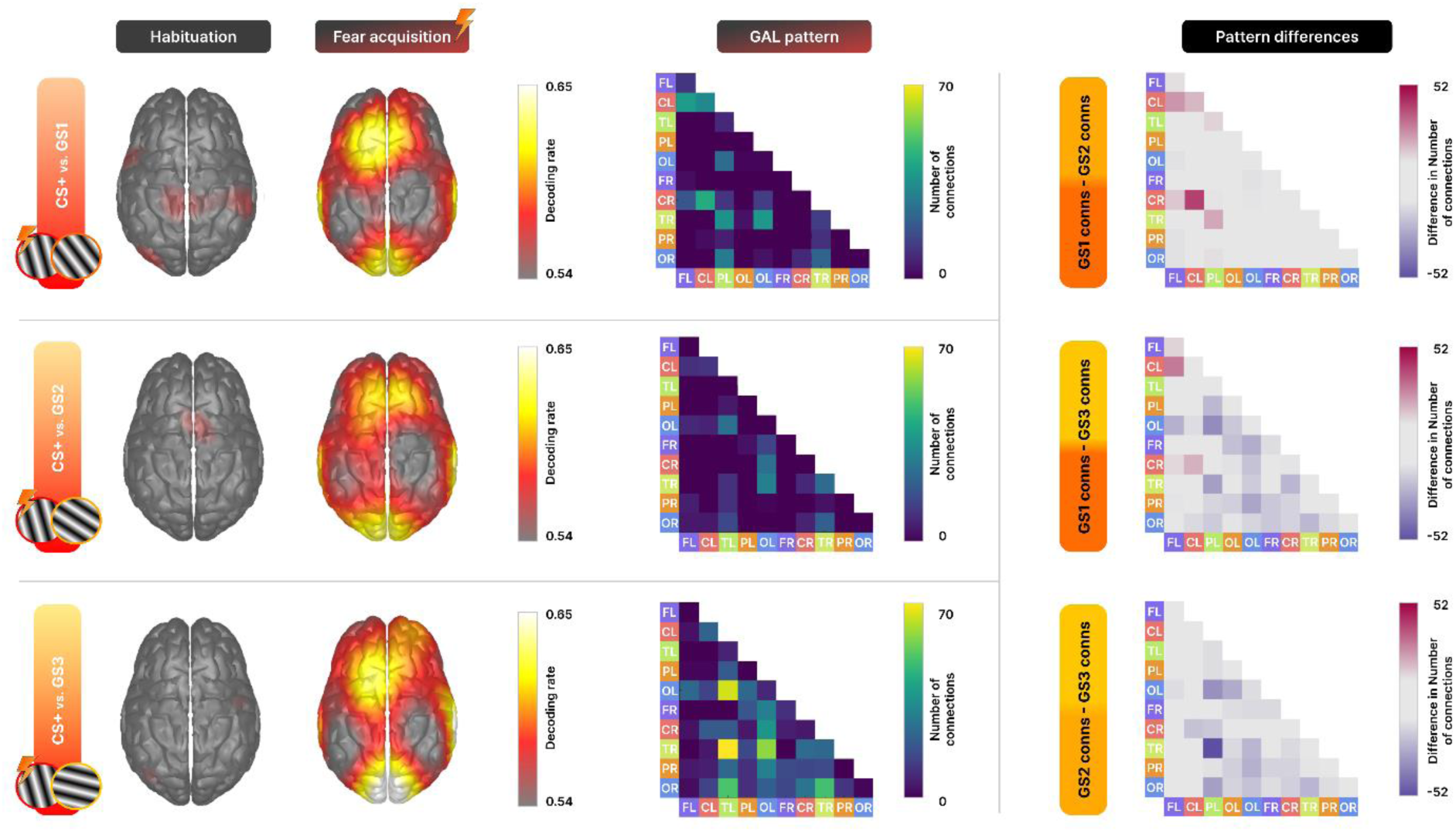
Changes in decoding and GAL connectivity patterns during fear acquisition. **Left**: Brain projection of decoding accuracy between CS+ and GS stimuli. The left brain refers to the habituation phase where CS+ stimulus was not yet paired with the aversive US, while the right brain corresponds with the fear acquisition phase where it acquired fear-related properties. **Middle**: GAL connectivity pattern for the three CS+ vs. GS pairwise based on the number of statistically significant generalization connections (Bonferroni-corrected α = 0.01) within brain areas during the fear acquisition phase. **Right**: Differences in generalization connectivity pattern between the GAL from each one of the three pairwise. Abbreviations stand for F = frontal, C = central, T = temporal, P = parietal, O = occipital and L or R for left and right, respectively.

## Discussion

The emergence of multivariate decoding techniques has made it possible to quantify and discriminate dynamic changes in neural patterns. Here, we present a decoding approach that uses the temporal patterns of neural population signals. The time-based decoder is then used to quantify the similarity of time courses across different locations. Furthermore, the combination of this decoding approach with a forward model visualizes the changing cross-decoding over time and thus illustrates the temporal sequence of spatial activation patterns.

The present study demonstrates the usage of the time-GAL method, employing two EEG datasets recorded during two tasks, both involving vision and emotion. In the first study, consistent with a large body of evidence [28], we found strong decoding during the period of the late positive potential (LPP) ranging from 0.5 s to 1 s post-stimulus. However, previous univariate studies rarely observed differences between pleasant and unpleasant content, supporting the notion that the multivariate time-GAL approach is more sensitive than univariate approaches. In addition, while univariate models [1] have occasionally identified condition differences between pleasant and unpleasant content during the LPP time window, the MVPA-based approach extracted the spatial pattern differences and highlighted brain areas with shared decodable information. Therefore, the time-GAL has substantial potential for defining the spatio-temporal dynamics specific to processing emotional scene content.

Application of time-GAL to the aversive conditioning dataset (data set 2) illustrated how the method discriminated between neural responses to threat versus safety, after conditioning. Prior to conditioning, no decoding was found for the same stimulus pairs, highlighting the sensitivity of the method to acquired stimulus properties such as conditioned threat [20]. In our example, dense anterior to posterior generalization connectivity patterns were found for the comparison of the fearful CS+ and the most similar GS condition, consistent with notions of sharpened orientation tuning as a function of experience.

The decoding methodology proposed in the present work aims to offer a new tool for the analysis and study of neurophysiological data. This approach leverages the rich temporal information contained in electrophysiological time series. Instead of using spatial information as traditionally carried out in fMRI [4,5] and M/EEG [9,10,26], here we utilize the temporal dimension as features to feed the decoding model. To our knowledge, MVPA-based decoding approaches have not yet been applied to time-based features. Therefore, the present tool may be the first step towards suitable methods for characterization neural activity patterns while taking advantage of their high temporal resolution.

Decoding approaches have been applied to neuroimaging data in a diverse range of procedures. Bae and Luck [26] have conducted time-resolved MVPA approaches using averages of trials in order to obtain better noise-ratio information and reduce computational resources. This approach obtains the evoked information of the trials, yet loses the induced information of each individual trial [12]. Following their approach, we computed a similar version of the GAL procedure making use of averaged groups of trials (see S1 Appendix). Overall similar information was found in the generalization connectivity matrices using averaged data compared with the single-trial based method proposed in this paper. Interestingly, no statistically significant inverse generalization connections were found when using trial averages, potentially indicating the usage of non-phase locked activity for single-trial based decoding.

The time-GAL approach can be easily applied using the toolbox available in https://github.com/csea-lab/time-GAL. No statistical assumptions are required to be met and both stationary and non-stationary signals can be used for decoding. Therefore, this procedure can be utilized to analyze a wide range of cognitive and affective neurophysiological studies. In the present study, a current source density transformation was applied to minimize effects of volume conduction in EEG data, but this step is not required.

The second step of the time GAL procedure offers information about how decodable information is generalized across the brain. The generalization connectivity maps obtained through the algorithm may be suitable to examine the configuration of dynamic cortical networks [29]. Traditionally, these networks have been studied conducting functional connectivity analysis using measures that reflect the statistical dependencies between two time series, such as the Pearson correlation coefficient or other metrics derived from general linear models [1]. This methodology is often applied to resting-state recordings [30] and tends to involve filtering of the data to highlight a given aspect of the neural time series. Here we extract decodable information from the waveform itself at the single trial level, using all available data.

In conclusion, time-GAL is a novel algorithm for examining neural dynamics based on decoding. At its core, it is non-parametric in nature because it is based on decoding accuracy measured by leave-one-out cross-validation. Thus, the method has substantial potential for application to wide range of spatio-temporal biological signals, especially signals characterized by rich temporal information.

## Acknowledgments

The authors would like to thank collaborators from the Department of Psychology and the J. Crayton Pruitt Family Department of Biomedical Engineering at the University of Florida for fruitful discussions and helpful feedback on the project.

---

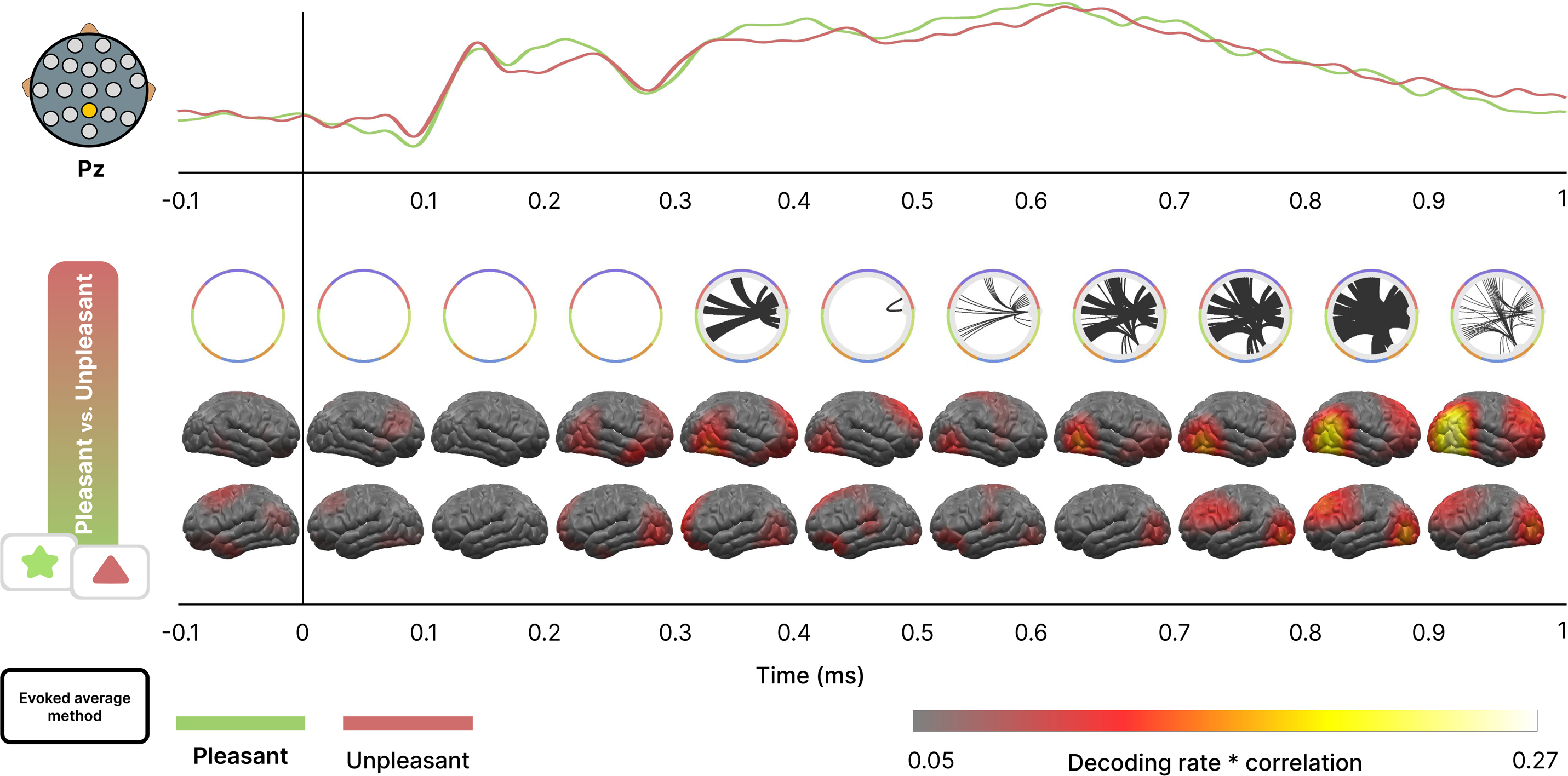

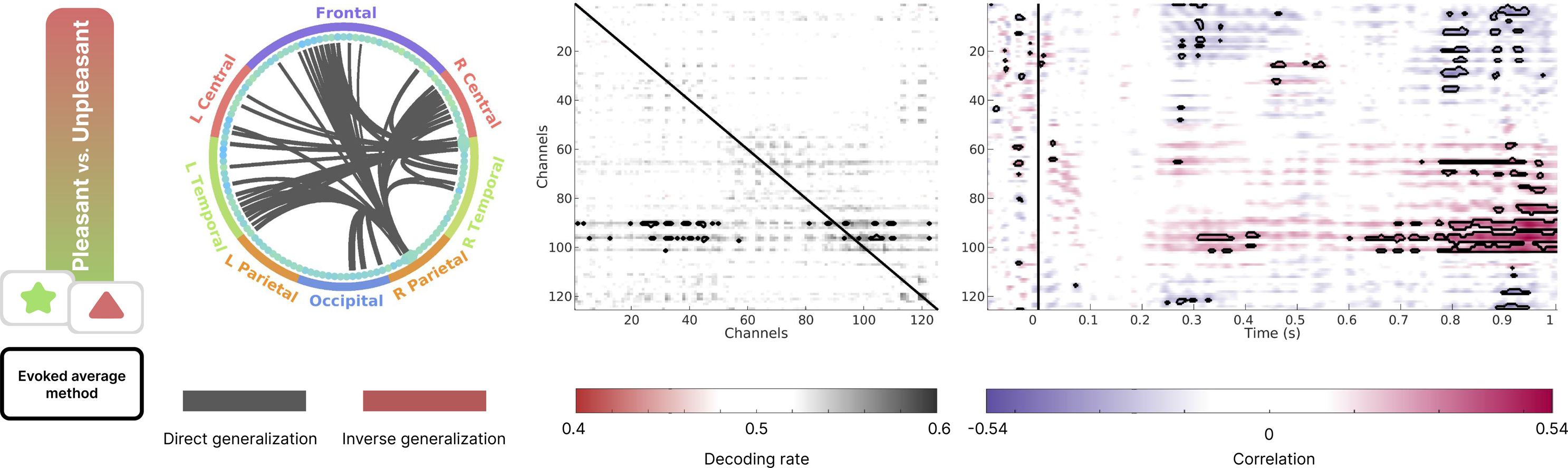

