## Appendix 1 for "Decoding in the fourth dimension: Classification of temporal patterns and their generalization across locations"

### Supporting Information 1 Appendix

Traditionally, the MVPA-based decoding approach has been conducted using spatial information to classify between conditions [1]. Usually, information from the experimental trials is condensed in averaged groups of trials to reduce the computational resources needed to train the model [2,3]. Therefore, classifier models requiring heavier computational resources, such as supported vector machine (SVM), can be used. However, in our work we employed linear discriminant analysis (LDA), a classifier method requiring less resources, thus allowing us to compute the generalization across location (GAL) methodology at the single-trial level.

Analysis of brain electrophysiological data following an evoked (averaging data) or induced (using single-trials) approach can derive different results [4,5]. For this reason, here we aimed to investigate whether the results of the GAL methodology using an evoked one similar as performed by Bae and Luck (2018) could differ from the induced approach developed in the main text. Particularly, in this complementary analysis we randomly divided the trials for each condition in three averaged groups using two for training the model and one for the test phase. Again, a leave one subject out cross validation was carried out training the model using *n*-1 subject and testing in the remaining subject. For simplicity and better comparison, ERP signal from the affective EEG task was employed since it benefits better from average than SSVEP signal.

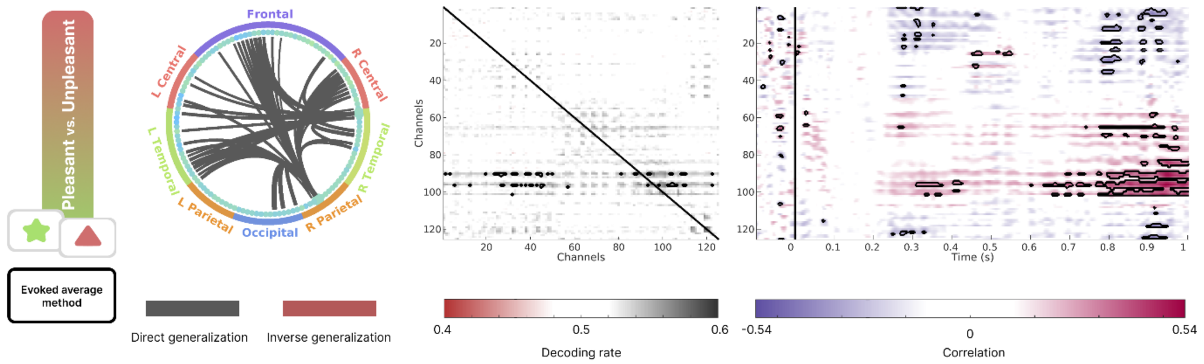

*Figure S1*. Generalization connectivity pattern and temporal distribution for affective pictures using an average-based evoked approach. **Left**: Circular graph showing the statistically significant generalization connections between 125 CSD channels. Black lines refer to those connections where both channels show similar decoding temporal patterns. **Center**: Generalization connectivity matrix showing decoding results (accuracy). Y axis corresponds with the channel data employed to train the model, while X axis refers to the channel data used in the test phase. Black contours indicate direct generalization connections after a Bonferroni-corrected statistical ($\alpha = 0.01$) comparison with chance level (0.5). **Right**: Temporal correlation between labels of condition and data showing the positive or negative weights of the decoding model. Black contours indicate those time points whose p-value is below 0.01 for each channel.

Interestingly, we can observe a lack of inverse generalization connectivity, indicating that the evoked method does not find statistically significant neural patterns of opposite response to the stimuli. In addition, induced approach found more contribution of the electrode's information to the discrimination of the conditions (Figure 3 right) than here the evoked analysis (Figure S1 right) from 300 ms to 500 ms. These differences may highlight the capability of the MVPA tools to extract the information from single-trials contained at the induced level and lost when information is averaged. However, late cognitive processing from 800 ms to 1 s shows similar contribution patterns in similar electrodes indicating that process could be more stable and less oscillation-dependent. Thus, difference between averaging approaches may not differ in late latencies. Regarding the generalization connectivity patterns (Figure 3 left and Figure S1 left), both analysis shows slight differences but preserving the same anterior-posterior generalization connectivity patterns.

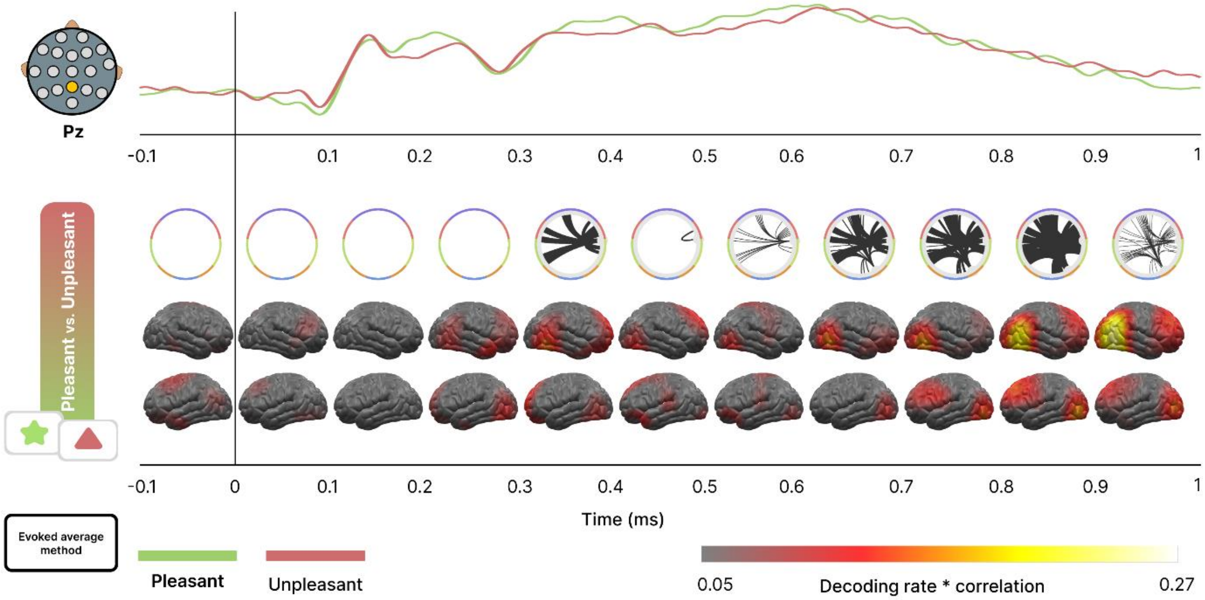


*Figure S2*. Time course of emotional content decoding and GAL connectivity patterns in affective pictures using an average-based evoked approach. **Top**: Electrophysiological ERP response to the presentation of different emotional content obtained at the parietal central (Pz) electrode for the three conditions (pleasant, neutral and unpleasant). **Middle**: Circular graphs show the GAL connectivity pattern at each time step for each pairwise condition. Brain's projections indicate the convolution of decoding rate by time correlation describing the fluctuation of the emotional process during the trial over the cortex.

Inspecting the decoding topographies (Figure S2 bottom), we can observe similar decoding patterns than those obtained using an induced methodology (Figure 4 bottom). Specifically, same anterior and posterior distribution of the neural patterns are found, overall, during the last part of the trial (800 ms to 1 s). This similarity shows that same areas and latencies are involved in the discrimination between pleasant and unpleasant conditions even when an induced or evoked methodologies are conducted, thus suggesting that both approaches are valid to analyze and extract conclusions from data.

In addition, GAL connectivity patterns shows their activity in the same latencies as previously indicated (Figure S2 top). In contrast with the induced methodology, less GAL activity can be found during the first part of the trial (300 ms to 500 ms) using the evoked approach. These results could highlight the communality of cognitive processes in late latencies discoverable from both induced and evoked approaches. In contrast, the induced methodology shows optimal to extract the information related to the oscillatory bands phase during the early latencies.
